# A Rapid Screening of Resistant Tomato Cultivars in Early Seedling Stage Against *Ralstonia solanacearum* Bacterial Wilt Under Soil and Nutrients Free Conditions

**DOI:** 10.1101/2020.03.16.993139

**Authors:** Niraj Singh

## Abstract

This work reports a rapid screening study of tomato *(Solanum lycopersicum)* cultivars resistant to *R. solanacearum* (F1C1) bacterial wilt in early seedling stage (6-7 days old, cotyledon leaf stage) under soil and nutrients free condition. This study was performed in 1.5-2.0 ml microfuge tube where root inoculation approach has been used during infection under gnotobiotic condition and got completed within 2 weeks. After continuous post inoculation observation of disease progression in all nine recruited tomato cultivar, Ayush was found to be the most resistant (~15% wilting) while the Ruby tomato cultivar was found to be the most susceptible (~95% wilting) against *R. solanacearum* bacterial wilt. This comparative wilting and survivability study among nine different tomato cultivar under same conditions indicated that early seedling stage of tomato may be a potent and helpful tool for an easy and rapid screening of resistant tomato cultivar against *R. solanacearum* bacterial wilt disease at large scale level in a very economical way in terms of time, space, labor and cost. Over and above The soil and nutrients free approach presented here may be helpful for the rapid selection and development of resistant tomato cultivar and new plant protection strategies against *R. solanacearum* bacterial wilt disease.

## 1. Introduction

*Ralstonia solanacearum* is a Gram (−ve), soil-borne, vascular plant pathogen which causes a lethal bacterial wilt disease in more than 450 monocot as well as dicot plant species from 54 different botanical families (Hayward, 1991; Elphinstone, 2005; Wicker *et al.*, 2007; Genin, 2010) such as many important crops *viz*. tomato, potato, brinjal, chili, peanut, pepper *etc.* (Gabriel *et al.*, 2006). *R. solanacearum* is also known as the causative agent of brown rot of potato, Moko disease of banana and Granville wilt in tobacco (Peeters *et al.*, 2013; Li *et al.*, 2014). In case of *R. solanacearum*, due to extensive genetic diversity amongst the bacterial strains of this pathogen, it is now considered as “*Ralstonia solanacearum* species complex (RSSC)” (Fegan & Prior, 2005). Due to exceptionally wide range of host, aggressiveness and adaptability, this wilt pathogen is causing huge damage to tomatoes and many other agriculturally important crops in subtropical, tropical and warm temperature geographical regions of the world (Pradhanang *et al.*, 2005). More than that, there is no effective way so far is available to deal with this pathogen and its associated disease.

In addition to its lethality in wide range of plant species, *R. solanacearum* have an extraordinary ability to survive in soil and water for many years in absence of suitable host as well as form latent infections within indigenous weeds that create difficult situation for eradication of this bacterial wilt pathogen (Hayward, 1991; Wenneker *et al.*, 1999). Over and above, this bacterial pathogen in recent past is treated as second rank in the table of the most economically important bacterial plant pathogens across the globe (Mansfield *et al.*, 2012). *R. solanacearum* causing a big threat to agriculture, emerging as a one of the most devastating pathogen that makes substantial loss to the production of tomato and other important crops (Peralta *et al.*, 2006). Tomato is one of the most demanded and a consumed vegetable across the world which is rich in vitamins, minerals and fiber contents (Peralta *et al.*, 2006). *R. solanacearum* bacterial wilt of tomato and in other crops is difficult to control. Yield losses due to this pathogen can rich up to 80-90% of tomato production depending on different factors such as cultivar, climate and soil type (Elphinstone, 2005). Yet, there is not a single approach which has been reported to be potent for the restriction of this pathogen and its associated disease. Some of the commonly used antibacterial agents such as copper, ampicillin, streptomycin and tetracycline have been found exhibiting low efficiency on suppression of *R. solanacearum* and its associated lethal wilt disease in the field. However, these chemicals are not environment friendly and expensive to apply in the field (Champoiseau & Momol, 2008). In addition to that, use of antagonistic bacteria or virulent mutants of *R. solanacearum* was an alternative biological control approach to bacterial wilt but the implementation and results obtained under controlled laboratory conditions were not promising in the field (Saddler, 2005).

In the absence of efficient strategy to control and eradication of *R. solanacearum*, the best possible way to overcome bacterial wilt may be to use the resistant or moderately resistant host cultivars (Champoiseau & Momol, 2008). Till date, most of the studies of screening of resistance cultivar against bacterial wilt have mainly conducted in *Arabidopsis thaliana* and *Medicago truncatula* (Huet, 2014).

In the present scenario, prevention and control of *R. solanacearum* bacterial wilt, resistant cultivars selection seems to be an environmental friendly and promising approach. Even many resistant rootstocks have been developed by different scientific groups to control the *R. solanacearum* wilt (Nakaho *et al.*, 2000). However, this lethal wilt disease has also been reported to be present in commercially available tomato cultivars generated by grafting technique (Nakaho *et al.*, 2000). The review of literature on *R. solanacearum* bacterial wilt disease has indicated that resistant tomato cultivar may be a good and promising experimental approach. Even few screening works on resistance tomato cultivar against *R. solanacearum* in soil grown and late seedling stage (25 days old) by artificial wounding way of inoculation has been reported (Kim *et al.*, 2016).

In this context of resistant cultivar, this study is reporting an easy and rapid way to screen resistant tomato cultivar against *R. solanacearum* bacterial wilt in early seedlings stage (6-7 days old, cotyledon leaf stage). During this study, *R. solanacearum* (F1C1), inoculated in early seedling stage of nine different tomato cultivars by root inoculation approach in soil and additional nutrients free condition, were used to evaluate the susceptibility of all these nine different cultivars against same *R. solanacearum* (F1C1) strain and their associated wilt disease. (Singh *et al.*, 2018a; Singh *et al.*, 2018b)(Singh *et al.*, 2018a; Singh *et al.*, 2018b)^18, 19^After pathogen inoculation till 8th days of post inoculation, pathogenicity, disease progression and wilting amongst all those nine tomato cultivars were observed in same conditions which were recorded and analyzed accordingly.

## 2. Material and Methods

### 2.1 Germination of tomato seedlings for pathogen inoculation

Nine different tomato cultivars viz. Durga, Rocky, Akhilesh, Classic, Nun 7610, Navya, Param, Ayush and Ruby which are commonly available to farmers in Assam, Northeast India were recruited for this study. Tomato seeds of all nine cultivars were pre-soaked in sterile distilled water for ~ 48h or two days. All seeds were washed twicely with sterilized distilled water before soaking in distilled water. This was followed by spreading the soaked seeds on sterilized wet tissue paper in nine separate plastic trays followed by germination in a growth chamber (Orbitek, Scigenics, India) maintained at 28°C, 75% relative humidity (RH), and 12 h photoperiod respectively (Singh *et al.*, 2018a)(Singh *et*.,2018b). Sterile distilled water was sprinkled at the regular interval till 6-7days to sustain the better way of germination and growth of tomato seedlings.

### 2.2 Bacterial strains, growth media and culture conditions

*R. solanacearum* and other non pathogenic bacterial strains such as *Pseudomonas putida, Bacillus subtilis* and *Escherichia coli* used in this work were previously reported (Singh *et al*, 2018). Growth medium used for wild type *R. solanacearum* F1C1 (Kumar *et al.*, 2013), TRS1016 (mCherry tagged) derivative strains of F1C1 (Singh *et al*.,2018, Monteiro et al, 2012) and *Pseudomonas putida* was in BG medium (Bacto glucose agar medium) (Boucher *et al.*, 1985). Incubation temperature for *R. solanacearum* strains and *P. putida* was 28°C. *B. subtilis* and *E. coli* strains were grown in LB medium (Bertani, 1951) at 37°C. 1.5% agar was added for solid medium if and when necessary (Singh *et al.*, 2018a).

### 2.3 Preparation of bacterial inoculum and root inoculation

Freshly grown colonies of *R. solanacearum* (F1C1) on BG-Agar medium were added to 50 ml volume of BG broth medium with the help of a sterile loop and placed in a shaking incubator (Orbitek, Scigenics, India) maintained at 28°C and 150 RPM for optimum growth. After 24 hours of incubation, fully grown bacterial cultures were centrifuged at 4000 RPM (3155*g*) and 4°C for 15 min. After centrifugation, supernatant was discarded and bacterial pellets were re-suspended in equal volume of sterile distilled water to obtain bacterial concentration of ~10^9^ CFUml^−1^. *P. putida* was also grown in BG broth similar to *R. solanacearum* at 28°C while *E. coli* and *B. subtilis* cultures grown in LB broth medium at 37°C in a shaking incubator maintained at 150 RPM. Inoculums of different bacteria were prepared by following exactly the same procedure used in case of *R. solanacearum* F1C1 as mentioned in (Singh *et al.*, 2018a).

### 2.4 Root inoculation of *R. solanacearum* in tomato seedlings

Around 30 ml of *R. solanacearum* F1C1 inoculum (~10^9^ CFUml^−1^) was taken in a sterile tube. From the germinated tomato seedling bed, 6 -7 days old (two leaf stage) tomato seedlings were picked one at a time. Further root of each seedling was dipped for 1-2 seconds in the bacterial suspension or inoculum (up to the root-shoot junction) followed by transfer of pathogen inoculated seedling into an empty 1.5 or 2.0 ml sterile microfuge tube (Singh *et al.*, 2018a). All nine tomato cultivars seedlings root were inoculated by same bacterial inoculum and by same procedure. After inoculation, each seedling of recruited tomato cultivars were transferred to the microfuge tube and exposed to air for ~5 minutes (Figure-4) before adding 1.0-1.5 ml of sterile distilled water to each tube. In these all experiments, a set of forty seedlings (N=40) were taken for each tomato cultivar infection and control was taken separately. Here two types of control sets were taken, water control and non pathogenic bacteria. In first type control, set of forty tomato seedlings were mock-inoculated with sterile distilled water for all nine cultivar separately and in second type control, 40 seedlings sets of tomato cultivars were inoculated with non pathogenic bacteria *P. putida*, *B. subtilis* and *E. coli* strains by following the same procedure as mentioned in above lines (Singh *et al.*, 2018a). All the inoculated tomato seedlings along with the control sets were transferred to a growth chamber maintained at 28°C, 75% RH and 12h of photoperiod. Infected and control triplicate sets of seedlings were analyzed for disease progression and wilting till 8^th^ day post-inoculation and findings were recorded accordingly.

### 2.5 *R. solanacearum* colonization study in tomato seedlings

Further in colonization study, mCherry labeled *R. solanacearum* strain TRS1016 was cultured in BG broth medium in presence of appropriate antibiotic. Grown cultures of strains were pelleted down by centrifugation and 10^9^ CFU/ml inoculums were prepared by same procedure described for *R. solanacearum* in above section. Bacterial inoculums were used for infection by root inoculation approach in 6-7 days old tomato seedlings as stated above. After third day of post inoculation, tomato seedlings were employed for colonization study and infected seedlings were surface sterilized by the method of Kumar *et al*., (2016) (Kumar *et al.*, 2017). After surface sterilization, seedlings were observed under the fluorescence microscopy (EVOS FL, Life technologies) for bacterial colonization study in 40X magnification adjusted in RFP filter.

## 3. Results

Seedling’s root of nine different tomato cultivars *viz.* Durga, Rocky, Akhilesh, Classic, Nun 7610, Navya, Param, Ayush and Ruby were inoculated with the same bacterial inoculum to evaluate the susceptibility of different tomato cultivar seedling against *R. solanacearum* bacterial wilt (Figure-1 & Figure-2). Regarding susceptibility and disease progression, comparative survival studies among these nine recruited tomato cultivars resulted in significant difference at early seedling stage under same conditions. After eight days of post inoculation, it was observed that amongst all these nine cultivars, highest number (~ 95%) of seedling death due to wilting was found to be occurred in Ruby tomato cultivar while least number (only ~ 15 %) of seedlings got wilted in Ayush cultivar in compression to control sets. Both types control such as water control as well as nonpathogenic bacterial infected tomato seedling control, have shown 100% survival. This screening study amongst the recruited seedlings of nine different tomato cultivars resulted that Ayush cultivar was found to be the most resistant one while Ruby cultivar as the most susceptible tomato cultivar in early seedling stage against the *R. solanacearum* bacterial wilt disease (Figure -2). In addition to that, it was also very interesting to observe that the rate of disease progression and susceptibility amongst the different cultivars was not found to be the same (Which is clearly indicated in Kaplan-Mayer survivality plot). This work, first time reporting the possibility of early seedling stage of tomato (6-7 days old, cotyledon leaf stage) may be used as a potent tool for easy and rapid primary screening of resistant tomato cultivar against *R. solanacearum* bacterial wilt in soil and nutrient free condition. Colonization of *R. solanacearum* in wilted and infected seedlings were confirmed by fluorescence microscopy (Figure-3). In case of infected wilted tomato seedlings, *R. solanacearum* colonization were observed in the form of red fluorescence in stem and root portions while in case of control, no red fluorescence or colonization of *R. solanacerum* were observed. In case of Ayush resistant cultivar, number of wilted seedlings have shown pathogen colonization in both stem and root portion while in case Ayush survivor, pathogen colonization was not observed in stem portion of cultivar seedlings.

## 4 Discussion

This rapid screening study of resistant cultivar under gnotobiotic condition has given a clear indication that even in early seedling stage, tomato cultivars have differential susceptibility towards *R. solanacearum* bacterial wilt. Resistance development against bacterial wilt in plant species are generally reported to be mediated by quantitative trait loci but performance of crops are also variable and influenced by surrounding environment and other associated factors (Wang *et al.*, 2013). In this connection, one of the potential factor might be plant microbiota/microbiome, although the proper and exact mechanistic role of plant associated microbes related to pathogen defense has not been established till date (Kwak *et al.*, 2018). In several earlier studies, plant associated microbes are well reported to have some crucial roles in host’s health and protection against plant disease (Berg *et al.*, 2015; Castrillo *et al.*, 2017). In many literatures, bacterial endophytes are also reported to confer protection against many pathogens and its associated diseases (Compant *et al.*, 2005; Gómez-Lama Cabanás *et al.*, 2014) including bacterial wilt of *R. solanacearum* in tomato as well as in other related crops (Tan *et al.*, 2011).To understand the differential pathogenicity and disease progression behavior of *R. solanacearum* in the seedlings of the nine different tomato cultivar, a further detail investigation is required. In this primary study of investigating the correlation of seedlings susceptibility with the presence of cultivable endophytic bacterial diversity (microbiome) inside the seedlings (unpublished work) has shown resistant cultivar Ayush to significantly exhibit a high number of anti-*Ralstonia* bacterial population in comparison to most susceptible Ruby cultivar. Further, this study had also investigated and observed metabolomics (HR-LCMS) study of seedlings of resistance Ayush and susceptible Durga ruby (unpublished work) which also had shown very significant variation in terms of metabolic profile. In future, in-depth study based on this primary observation may contribute for better understanding of host-pathogen-endophyte interactions and its role in host resistance development in tomato against *R. solanacearum* wilt.

## 5 Conclusion

This screening study results has concluded that Ayush was found to be the most resistant amongst nine employed tomato cultivars. Over and above, it also indicated that early seedling stage of tomato (6-7 days old, cotyledon leaf stage) may be a potent tool for easy and rapid primary screening of resistant tomato cultivar against *R. solanacearum* bacterial wilt at large scale level in a very economical way in terms of time, space, labor, cost. It can be very helpful for scientific communities and reputed seed companies for resistant cultivar selection and its associated research.

## 6. Acknowledgment

Niraj Singh is very much thankful to Tezpur University, Assam, India and Royal Global University, Assam, India for providing an opportunity to work. Over and above, Niraj Singh is also thankful to Prof. Suvendra Kumar Ray, Tezpur University, for motivation and kind support. In addition to that, Niraj Singh also thankful to DBT,Government of India for the DBT fellowship and providing necessary facilities.

---

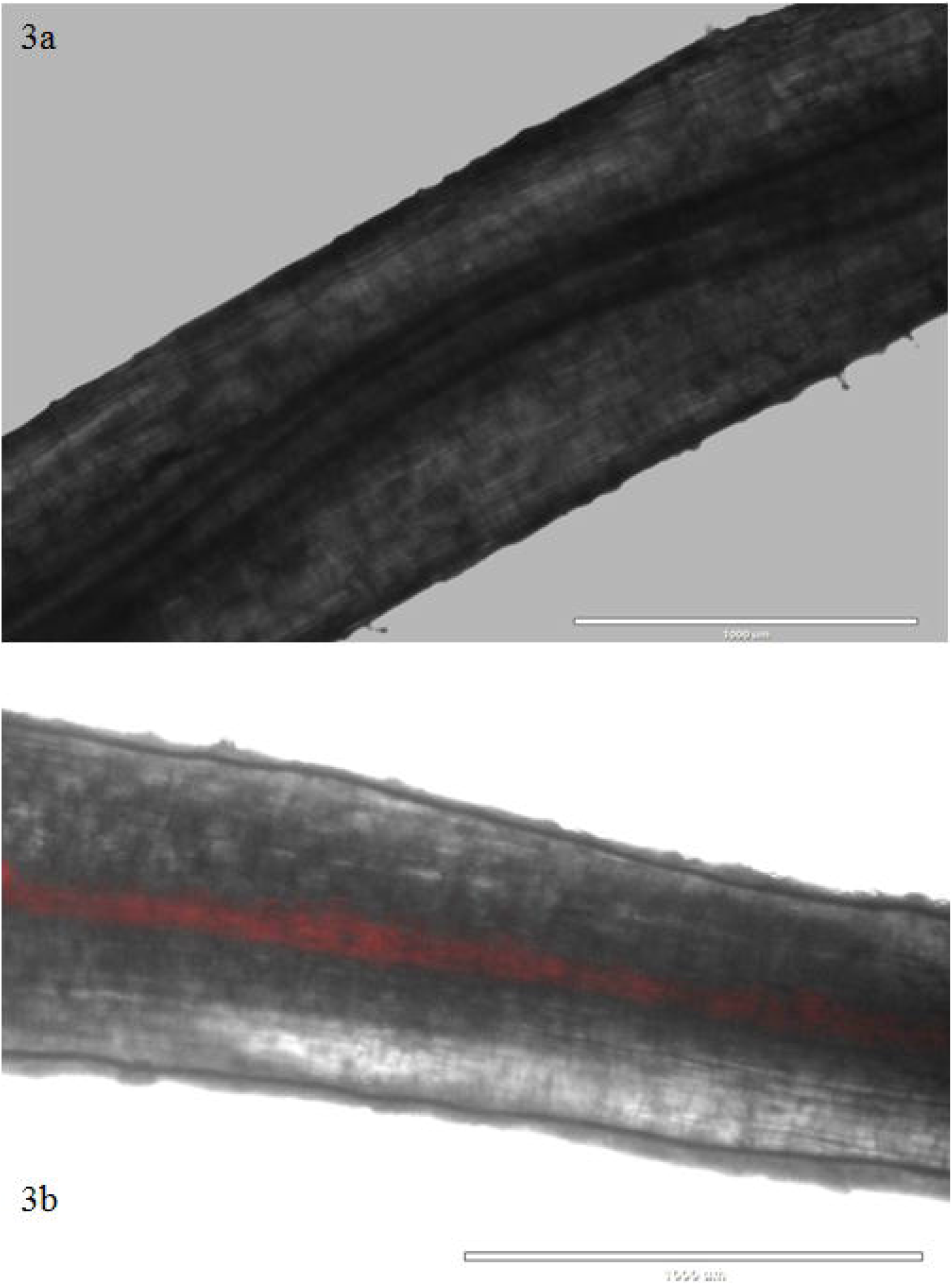

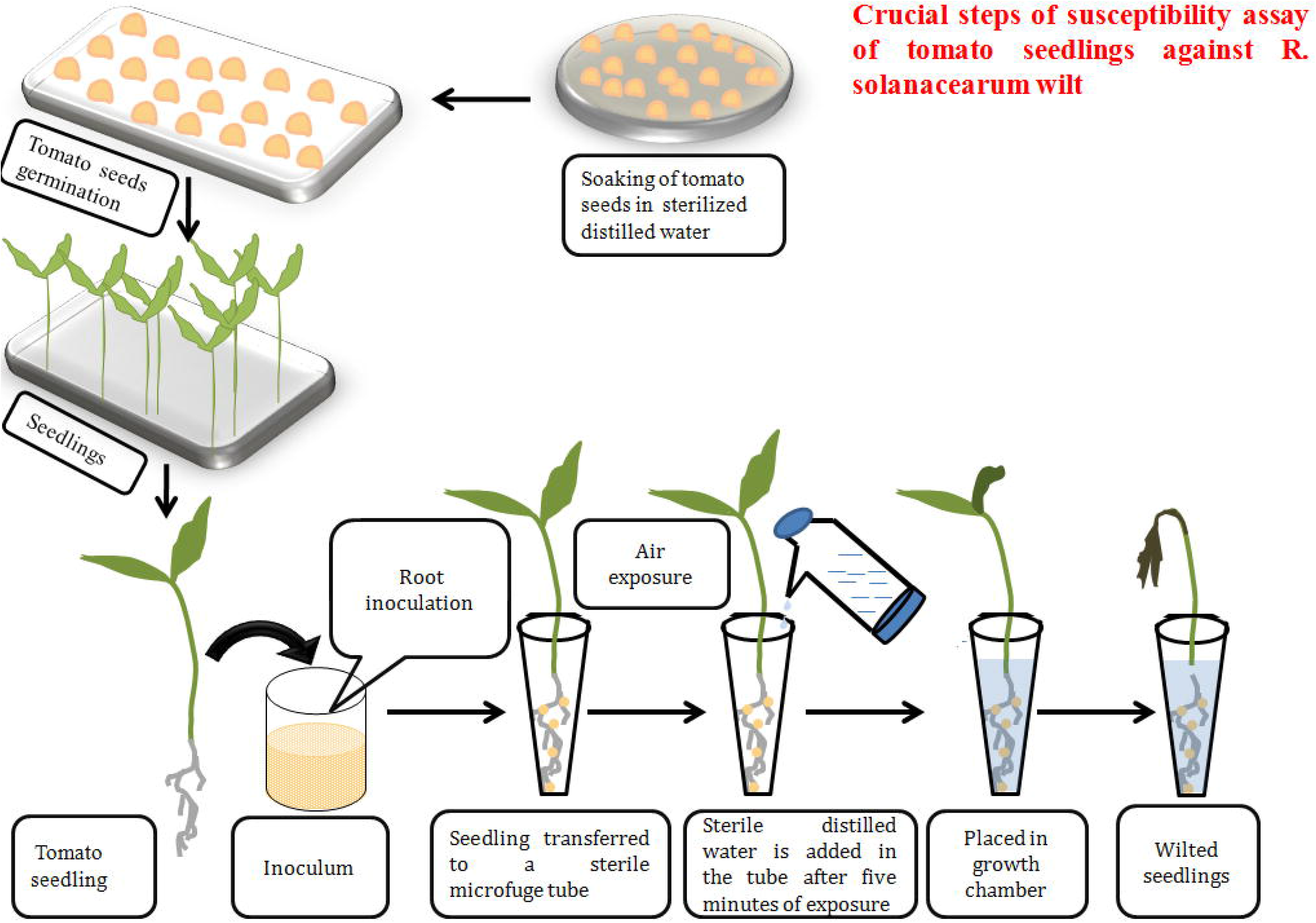

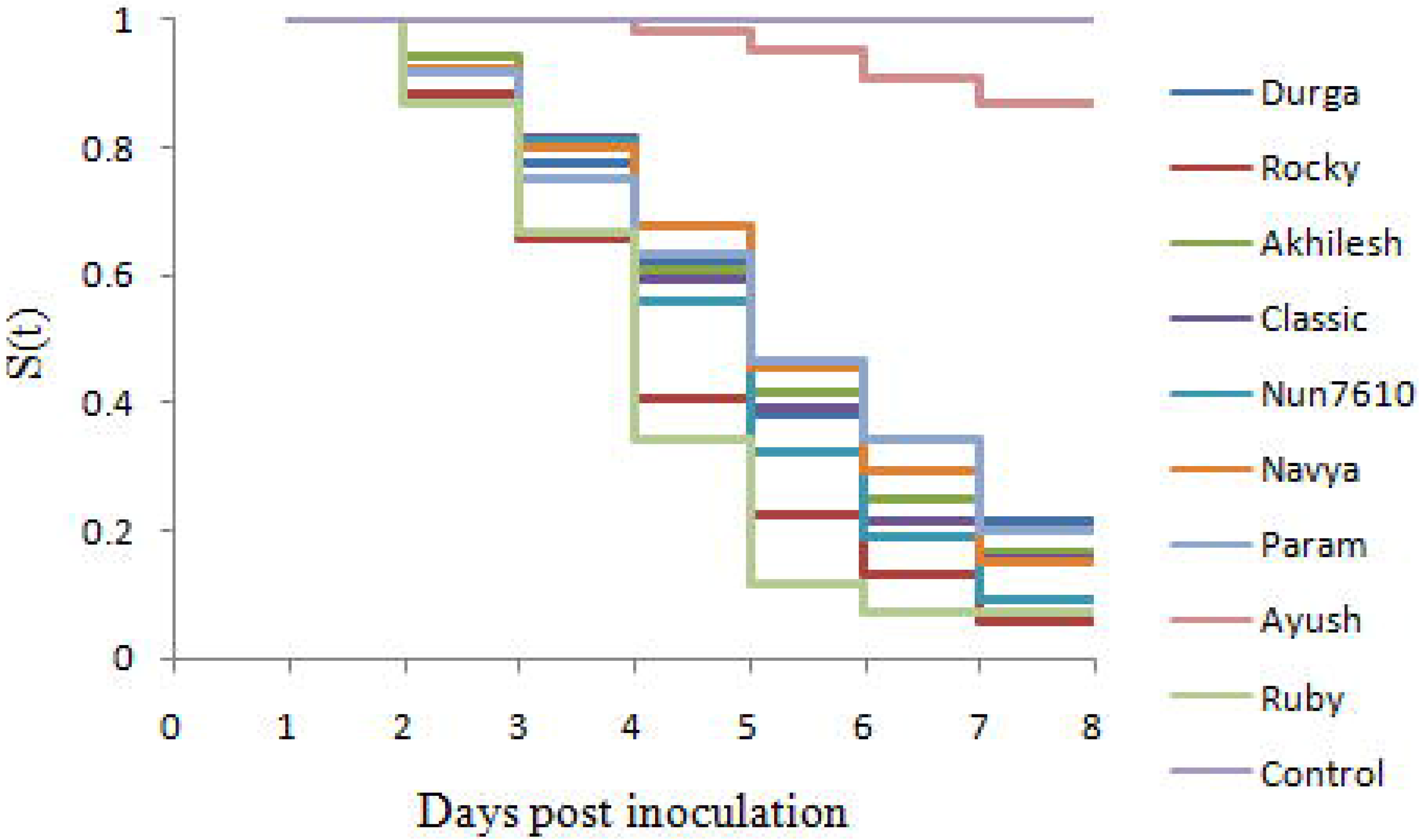

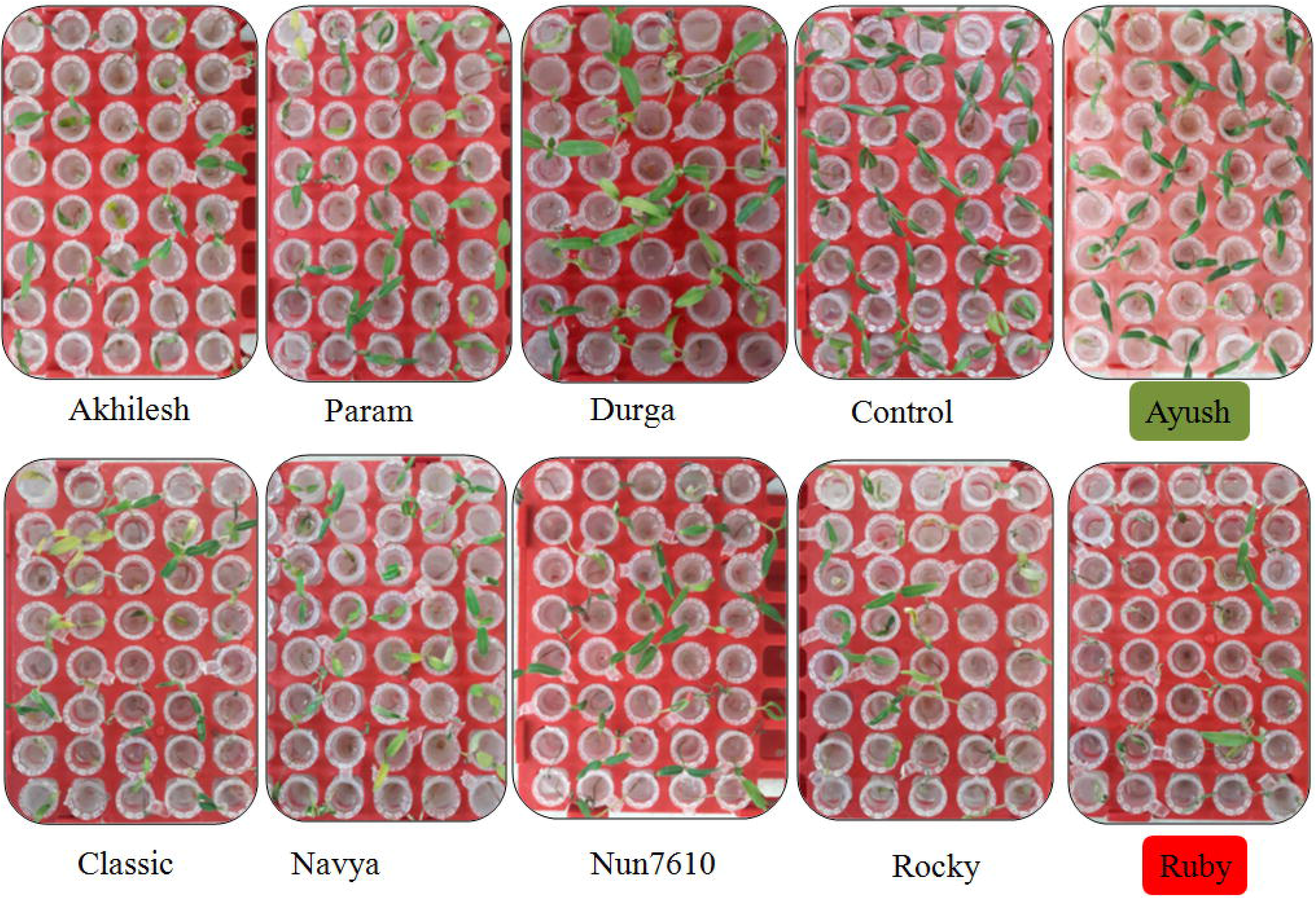

